# Study on tumor cell RNA cross-anticancer mechanism in non-tumor mice experimental model

**DOI:** 10.1101/277228

**Authors:** Qiwang Xu

## Abstract

To report the experiments of fractal structure and self-similar function in biological coupled oscillation brings life system waving growth in life organism. At the same time by self-organization the system openness come into being to maintain stability features of growth survival. From this, the orderly oscillation of tumor cell RNA was recognized that this process could lead to system sequentiality openness. According to waving growth of basic research the procedure could be sure to turn into a model as out of control of growing in tumor. Then in complete contrast that derived from the same component which have cross resistance effect. It shows that its biological behavior is converted as only a single physical power to play action by inactivated processing. Subsequent further confirms the essential roles whether essential roles link up with recovery of physiological function. The result in that it could exhibit feature of anticancer owing to be endowed tissue embryonization response. So that it also exerts preventing on senile chronic diseases. In theory, the concept of this tissue response in life organism closely link up with the single physical power of RNA component so as to bring up specific procedure of growing and subsisting that deal with keeping life vigor generalized explanatory.

## INTRODUCTION

The reason why cancer is called an “incurable disease” is due to lack of fundamental understanding of oncogenesis [1–2]. Over the past forty years on both the scale and the scope of scientific research on cancer had reached an unprecedented level, with the emphasis on cancer vaccine [3], new drug and research on new medical technology. It has constructed a new science in contemporary medicine. The studies on biological wave started since the second half of 20th century [4]. The experiments at earlier stage were based on random operations, by using lysogeny broth (LB) medium and adding 2, 3, 5-triphenyl tetrazolium chloride (TTC) as a growing reaction reagent, to observe swarming growth of proteus mirabilis via biological coloration as shown in Fig.1. The same kind of bacteria appearance as two different colonies was regarded as phase variation phenomenon in the process of bacterial growing. The result caused by adopting connecting culture technique has shown that is a process of limitless growing. Till the basis of single bacillus population biological wave mathematical model established by Jiuping Xu et al [5–6], the compound noun “biowave” of biological wave started to be used. From then on, the research on human biowave testing technology [7] has been set up. The neutrophilic granulocyte in blood was selected to do the test targets, using its characteristic of being sensitive to represent its integral change in real time. This sensitivity can also be shown as an enzymatic reaction to the redox of individual cell. The test used (3-(4,5-dimethy thiazolyl-2)-2,4-diphenyl tetrazolium bromide) (MTT) as hydrogen acceptor and formed colored formazan in the cell, so as to count the cellular rate in the positive or negative reaction. It represents the strength of integral cell population redox reaction. The study of biowave emphasizes the role of cell membrane permeability to the biochemical reaction of oxygen intake. In living organism the cell population had been measured to determining the base proportion changes at different times. The quantitatively elucidated ratio of redox changing dynamic process is termed “human biowave” dynamics [8]. The random change of biowave dynamic state could reflect the uncertainty characteristics in biology. Thus the chaotic concept of life organism was introduced into the research field at that time. This kind of uncertainty of life organism resulted in biowave effect time uncompressing technique arose. By the first time, it made human experimental forecast come true. Human biowave experimental forecast research results [9] showed that the occurrence or change of tumor is accompanied by biowave dynamic corresponding change. Thus it would establish technical conditions for studying the link between cancer and life organisms. For this reason, the research was converted from biowave basic theory to build the “non-tumor mouse” experimental model. To further clarify the stability change of biowave in inoculated tumor cells before and after. This will create conditions for further understanding of the life organism’s response to different tumor cells and tumor cell components of experimental operation processing. This is good for clinical exploration of cancer. In systematic study, the result astonish that there is not only the growth of the tumor between the life organism and the cancer cells correlation, but will also the life organism can produce the anti-cancer ability. The research on the combination of biowave basic experiments and non-tumor mouse model shows that certain ingredients of tumor cells can also emerge anti-cancer promotion ability. And even more amazing is cross-resistance can occur among different types of tumor cell lines. As the research further develops this nonspecific effect property that directly point to a new field arises. Basic experiment makes clear that the efficacy of anticancer mainly rests with the potential effect of tumor cell RNA constituent. In view of the RNA constituent special preparation as an inactivated RNA constituent result in pathogenic ability of malignant proliferation is lost. So it tentatively is called as “inactivated RNA vaccine” or the RNA for short. According to the basic experimental analysis, the basic mechanism be identified that the key factor is from the aid of life organism vitality. Thus,the RNA could keep happening coupled oscillation sequence characteristic. Subsequently the system is in the negative entropy flow dynamic process. As thus it can quantitatively express the effect of the RNA in the same way. So it can have been verified by life organism biowave effect time uncompressing assay in cancer incidence investigation. It has been revealed the RNA possesses the forcefully the organism biowave regulating effect that maintains the biowave stable and strong dynamic tendency. According to the basic experiment analysis suggests that this effect rests with the coupled oscillation dynamic process of molecule structure. The importance of this result is to promote system openness in life organism. Analysis identification that the condition of openness system can not only eliminate the carcinogenesis ecological niche, but also prevent tumor recurrence in situ as well as distant metastasis after surgical removal. And even the more important is the RNA could strengthen physical vitality for the advanced cancer patient. So much that could improve the endurance in the process of surgery. In further explored the action mechanism of the RNA, the result had involved the RNA efficacy originated from the biological behavior of the inactivated RNA structure as single physical properties. It is this has been converted characteristic that could be possible to have such a special biological nature. Compared to the traditional basic medicaments the tumor cell RNA gives evidence the omnibearing efficacy in whole body. The effective fast and persistent as well as more targeted to prevent and control cancer.

**Fig.1.**
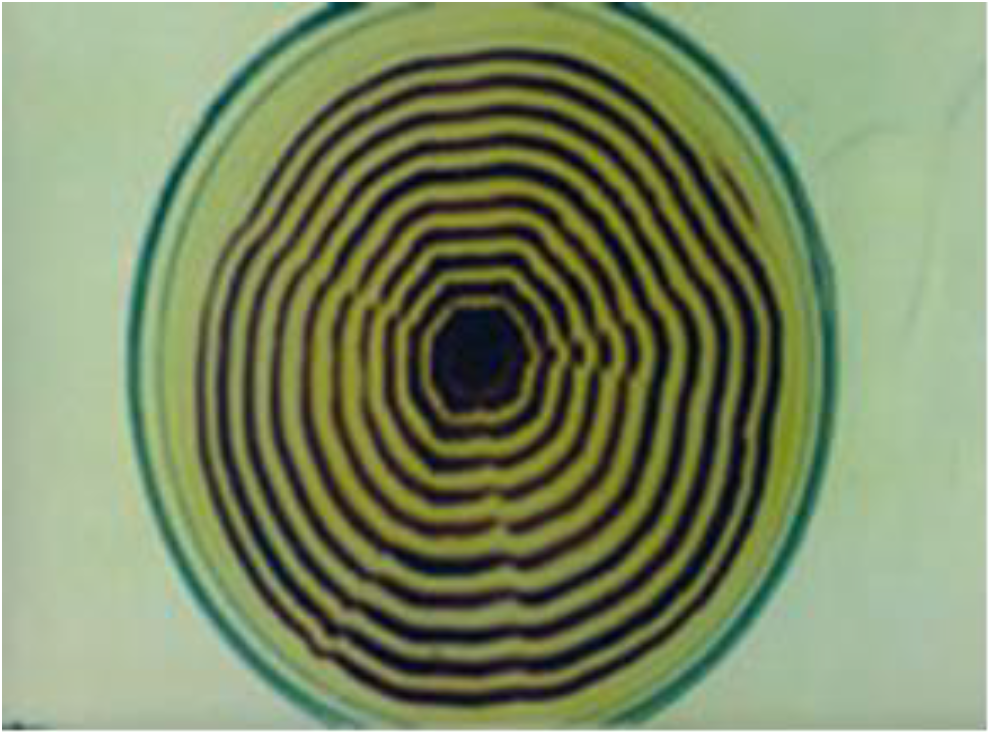
the dynamic biowave growth.

## BASIC EXPERIMENT AND RESULT ANALYSIS

### 1. Biowave Experiments

#### 1.1 Waving Growth Culture

Adopt LB solid medium add 0.05% TTC. Point—inoculate the strain onto a center of the plates. The incubator conditions have been regulated as temperature of 32 – 39 °C, relative humidity 40%, and keeping ventilation.

#### 1.2 Waving growth

The red bacterial colony is called central colony, came into being in 8 hours. Later on, yellow concentric ring bacterial colony encircles the red central colony formed. Thereby, it began to start the alternative waving growth process of yellow and red concentric ring bacterial colony every three hours, as shown in Fig.1.

As picture above showed that in the bacterial phase variation, the red colony is the vegetative cell and the yellow is the cryptic growth cell. The red central colony is the basis of waving growth. The size and shape of its colonies decide the subsequent dynamic condition of waving growth ring. It reflects the “starting control effect” of waving growth and shows the self-similarity characteristic in the growth system. In the process it formed a natural system that the vegetative cell proliferation and metabolism. Then the cryptic growth cell strenuous movement to keep structure away from balance. Result in the life organic tissue always as negative entropy flow sequentiality dynamic. The waving growth only ends until it reaches the rim of LB medium, which shows the finite limitlessness of waving growth. Further adopts the “connecting culture technique”. The results displayed the waving growth still keeps on though it had reached the length of 7.2 meters which verified the continuous growth characteristic. Currently experimental studies have formed a definitely reasoning that the tumor cell RNA component showing sequentiality dynamic effect always occurred imitating the bacterial wave characteristics.

#### 1.3 The periodic change of wave growing traces

The process of bacterial waving growth made the agar medium corresponding to concentric ring colonies alter. Uses cresol red indicator color reaction, and thus the medium under red colony were red. With the appearance of yellow colony, the color of indicator fades away gradually, as shown in Fig.2. One kind of bacteria species can grow out two kind of colonies with corresponding growth traces chromogenic reaction which shown the periodic characteristic in the process of waving growth. Combining with the distinct appearances of the micro-environments of colony related medium, it has further defined the biological phase variation process from red ring to yellow ring. So it is accompanying with the change of biological basic feature called as the phase variation. Before it the bacterium is in vegetative phase as basic form, while after phase variation the bacteria appearing slender shape also known as filaments peritrichous flagellum called as bacterial “cryptic growth cell”. It has active movement and the biochemical metabolism had changed accordingly at the same time, as shown in Fig.2.

**Fig.2.**
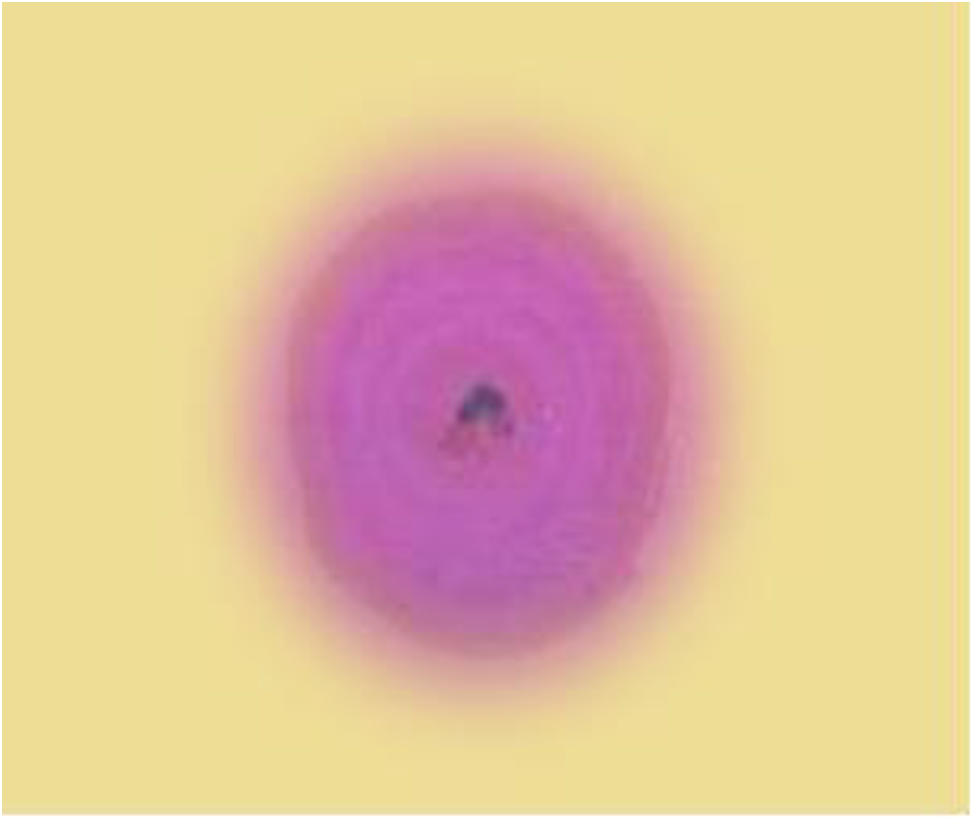
the medium of cresol red indicator coloration.

Fig.2. Showing above the medium of cresol red indicator color reaction. Its corresponding medium of “cryptic growth cell” yellow colony represented no metabolic waste products. During phase variation process exhibits the periodic alteration of waving growth. Not only “cryptic growth cell” do not undergo the usual biological metabolism but also it prompts the dispersion of metabolic wastes. It constituted an open regulatory mechanism of the growth system to periodically maintain open basis for the life organisms. It is compounding colonies express the process of culture and growth. The medium of the central red colonies appears in red color. The color of red rim gradually fades, and then goes away appearing without color, while pH of the agar medium in the yellow colonies has not changed as seem no biological growth even the medium under the same conditions. Then the two kind of coloration was the same bacterial diverse timing reactions.

#### 1.4 Biowave makes limitless growth

Through bacterial waving growth as description in Fig.1 and 2 reveal the basic theoretical mechanism for limitless growth. It is speculated that tumor cell RNA vaccine effect also as the same. It’s just the biological effect of non-life characteristics only. The red is basic growth, then the yellow colony consists of the cryptic growth cell of slender shape with moving and non-growing. It takes place by turn between the two forms continuous growth in order to maintain the advantage of quantity of species. Among phase variation process the cryptic growth cell play an important role. It proves that the pause of usual life activities is an indispensable step in sustaining survival. So the preventive measure of resting strain is a universal law in nature biology.

#### 1.5 Waving rejection experiment

There was the fast and slow cell line of dual phase cell line in phase variation. They are the quick or slow strains. The quick waving strain was inoculated in the medium center. The slow waving strain was inoculated the predicted fifth position of waving growth system. Nonsynchronous waving growth of theirs has been observed. Slow waving fan-like surface forms narrow-interval bacterial colony, as shown in Fig.3. The bacterium growth represents two different kinds of waving speed, and the interval between each waving ring remains unchanged. This bacterium which represents different waving speed grows apart, and it can also maintain its own growth characteristics.

**Fig.3.**
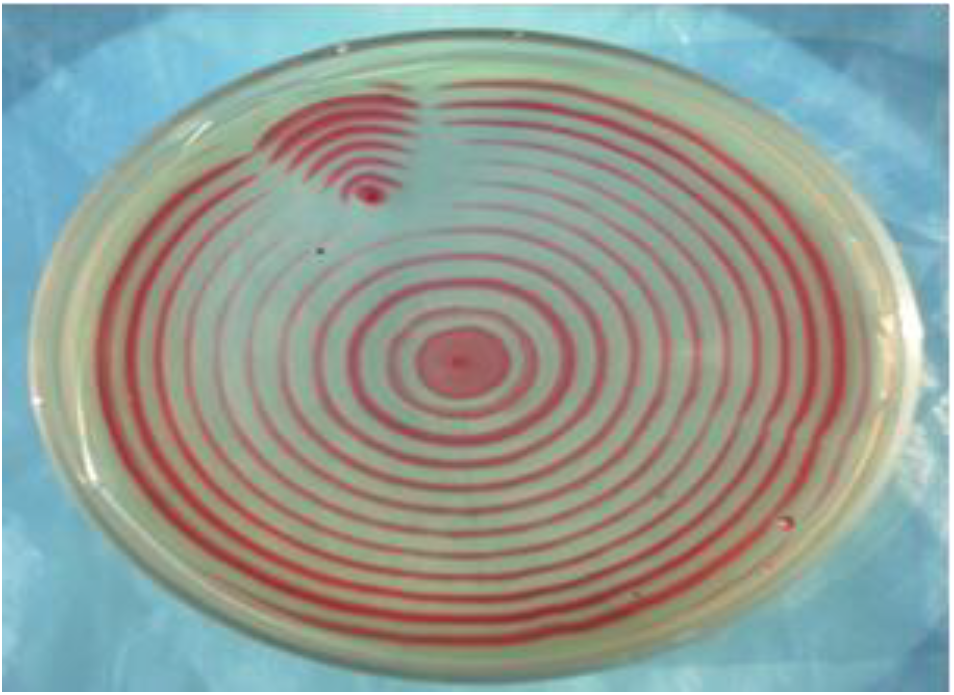
Quick-slow waving growth non-fusion phenomenon.

The Fig.3 above showing the culture result of quick-slow waving growth. One is the quick waving growing concentric ring colonies which are pointed in the center; the other is the fan-like colonies which are located on the fifth position. The interval between both of the waving rings has obvious differences but meanwhile remains unchanged.

In the subsequent experiment [10], quick and slow waving growing strains were selected and the two kinds of bacteria were mixed with the proportion of 73:100, followed by making the mix strains point inoculation and cultivating their waving growths. The specimen from first to sixth concentric ring colony was collected from each of them, and the dynamic change of the two bacterial quantity ratios in the process of waving growth was analyzed using the qualitative and quantitative culture technique. The results showed the proportion of quick waving bacteria decreases obviously on the first ring colony, which is 30.4%; the second ring is 16.8%; the third ring is 9.3%. The quick waving bacteria cannot be detection on the fourth ring, and it only shows slow waving bacterial growth. The result of quantitative analysis experiment has shown that bacteria which have mixed and grown with others as the different waving speed represent rejection biological characteristics. The experimental result has also shown that, with the consistency of its waving growth, life organism can get the power of “bio-dynamic selection”. Under the dynamic environment of waving growing and surviving in the phase variation process, the “cryptic growth cells” that the quick waving cells have transferred earlier, so that do not adapt to the environment. Therefor they suffer from repulsion in the ecologic competition. The biology kinetic energy is shown as waving growth repulsion represents the power of synchronous biology phase variation. It can prevent the tendency of disproportionation development from heterology changing cells in different directions. Meanwhile, this process lays the system periodically. It should be mentioned that the reaction of waving growth repulsion can directly display a screening mechanism to cell population in all kinds of tissues. Biowave research analysis regard as it’s an identification and selection process for the effect of tissue embryonization reviving plasticity. On this account, the biowave principle of the species stability has been verified. The following experiments further clarify the screening power of life tissue.

### 2. Experimental observation of cryptic growth cell

#### 2.1 Single cryptic growth cell movement

In single cryptic growth cell movement is the essential characteristic. It shows the slender cryptic growth cell and its essential characteristic of single cryptic growth cell is moving as the following:

**Fig.4.**
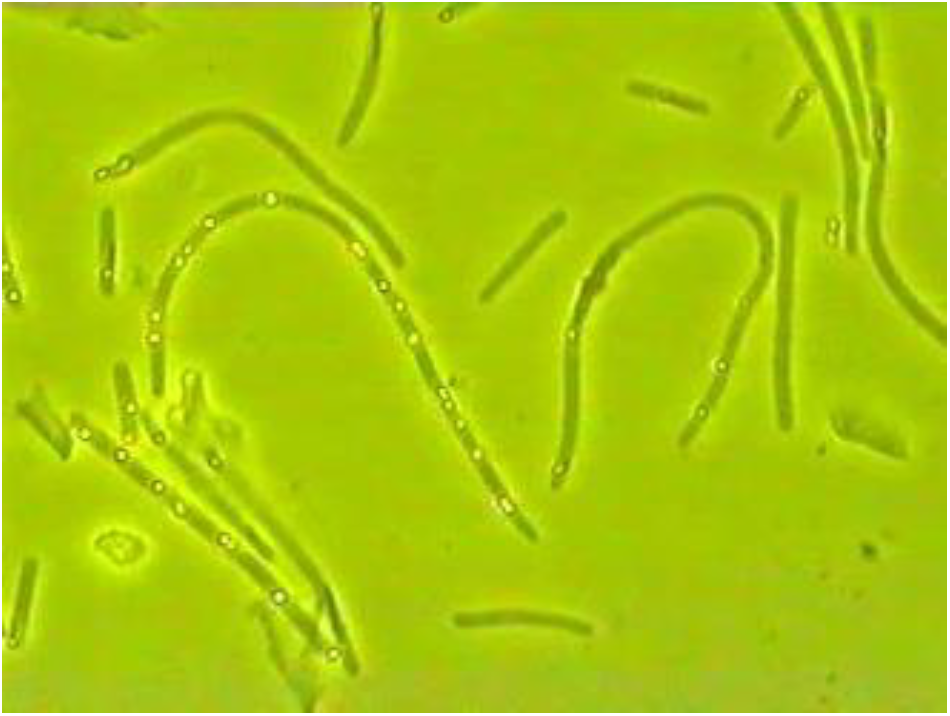
The basic feature of single “cryptic growth cell” movement.

It shows that the slender shape of cell has the ability of supporting movement and can remove all barriers. This means the strong features of power in life organism even micro-structures. Meanwhile, the single slender cell has shown the active movement characteristics of the randomness in the curvature movement. It showed that the cells would be as in the form of cryptic growth cell that is to maintain the power of movement through non-consuming processes. The results confirm that the power is derived from another approach. To this end, the fractal structure of coupled oscillations and its function are introduced.

#### 2.2 Bacterial cryptic growth cell coupled oscillation experiment

The cryptic growth cell incubation experiment referred to J. A. Shapiro method [11] of the microscopy-observed bacterial growth dynamic movement as the basis. Used the Opton El-Elinsatz phase difference microscope observed the cryptic growth cells display unusual active movement. That can prompt the occurrence of coupled oscillation as one process of term of physics had shown in Fig.5.

**Fig.5.**
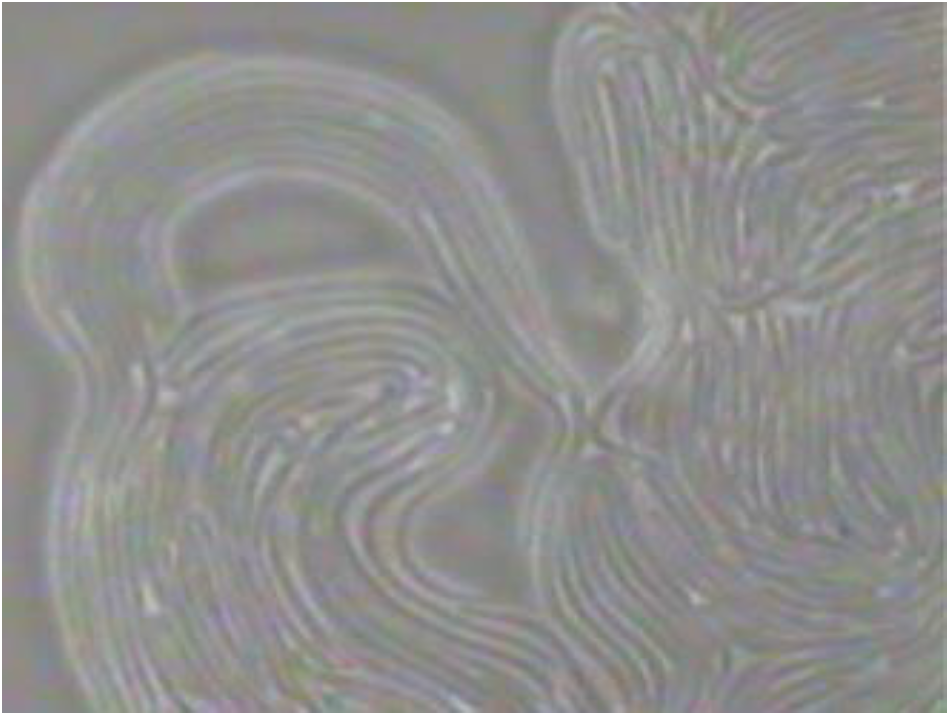
“cryptic growth cell” coupled oscillation feature.

When observing it with the microscope at 1000 times magnification, this kind of slender shape group will form into the process of coupled oscillation. The slender cell population can form the synchronous active orienting movement. When the moving condition is permitted it can keep on moving ceaselessly. Thus, the bacterium as cell population can form coupled oscillation. So-called cell is the basic unit it also should be the basis of dynamic structure of biowave. The bacterial cryptic growth cells can constitute coupled oscillation dynamic process generates a tremendous of power to promote the biochemical metabolic waste far away from the growing tissues. So that it exhibits survival capacity and reveals the limitless vitality in life organism at once. From this, there is corollary that the coupled oscillation process is exactly what the mechanism of RNA component display roles as a Nonliving material structure. So as to the coupling properties play a lasting effect with the aid of the dependent cell vitality. This is the theoretical foundation of the RNA vaccine efficacy.

#### 2.3 Cryptic growth cells subcellular structure coupled oscillation

The bacterial subcellular structure, flagella coupled oscillation is special phenomenon. In addition to bacterial “cryptic growth cells” could arise couple oscillation. The bacterial flagella as subcellular structure also produce similar forms of motion showing the self-similarity characteristics. So it called as fractal structure of bacterium. Under microscope the waving culture growing rim position of the yellow ring colonies had been observed that multi-strip cryptic growth cells can be seen the ordered side by side under the oil immersion lens. The synchronous sequence oscillation occurs among the flagella on the surface of each “cryptic growth cell”. Under the microscope It shows the kinetic characteristic of subcellular structure coupled oscillation and multiple oscillating shade-like chains consist of dynamic schistose shape, which seems like the cloud drifting phenomenon as shown in Fig.6.

**Fig.6.**
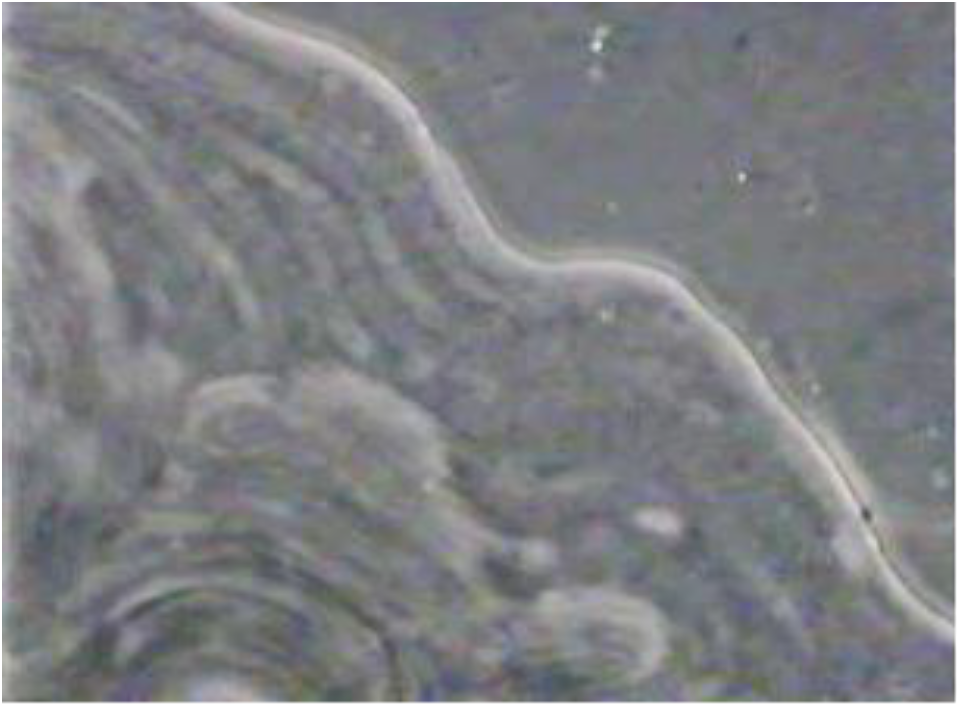
Bacterial subcellular flagellum sequentiality coupling oscillation.

Bacterial growth cell subcellular coupled oscillation expresses two self-similar oscillating systems. It revealed that biological organisms are likely to have a general oscillation system from structure to function. In this way the systems of life organisms could show the survival stability with the coupled oscillation tends to the sequentiality dynamic randomness. In living organism this feature maybe with universality and objectivity, so biowave research introduces concepts of biological coupled oscillation, subcellular structure coupled oscillation, biological coupled oscillation fractal structure that shows the different levels sequence oscillation in a same cell structure. And the nonlinear relationship sets up among the similar oscillatory structures in a cell.

#### 2.4 Cryptic growth cells perform biospin

The stability movement of “cryptic growth cell” lies in its form of spin which coiled multiple ring structure. This structure is called bio-spin. It can maintain the endless rotations till the life condition vanish. Single or multiple bio-spins can be observed in a visual field under the microscope. The rotation direction of theirs differs from each other. Its rotation velocity is basically consistent, showing the stable supporting the dynamic activity power. Fig.7 shows the fundamental feature of bio-spins. Its rotation as the consistency of the speed shows the stability of power while the difference of rotation direction reflects randomness of movement. These contribute to form waving motion which is to favor produce liquid convection. From the integral sequentially and non-repeatability, namely for the liquid in the endless turbulence have laid a foundation. The experiment followed J. W. Costerton called the bacterial cryptic growth cell act as the bacterial filaments [12]. Biowave experiment selected the edging of waving colonies under microscope observed the filaments motility phenomenon. The most special was the filament could spontaneously generate rotating simple structure called as bio-spin. It is very high rotational stability so as to almost never stopped that shows the limitless source of power. In terms of strength, its ability is more lie in the non-uniformity of size of bio-spin. Non homogeneous individuals cluster in essence, it Also belongs to group self-similarity. Detailed analysis Fig.7, especially combined observations PPT

**Fig.7.**
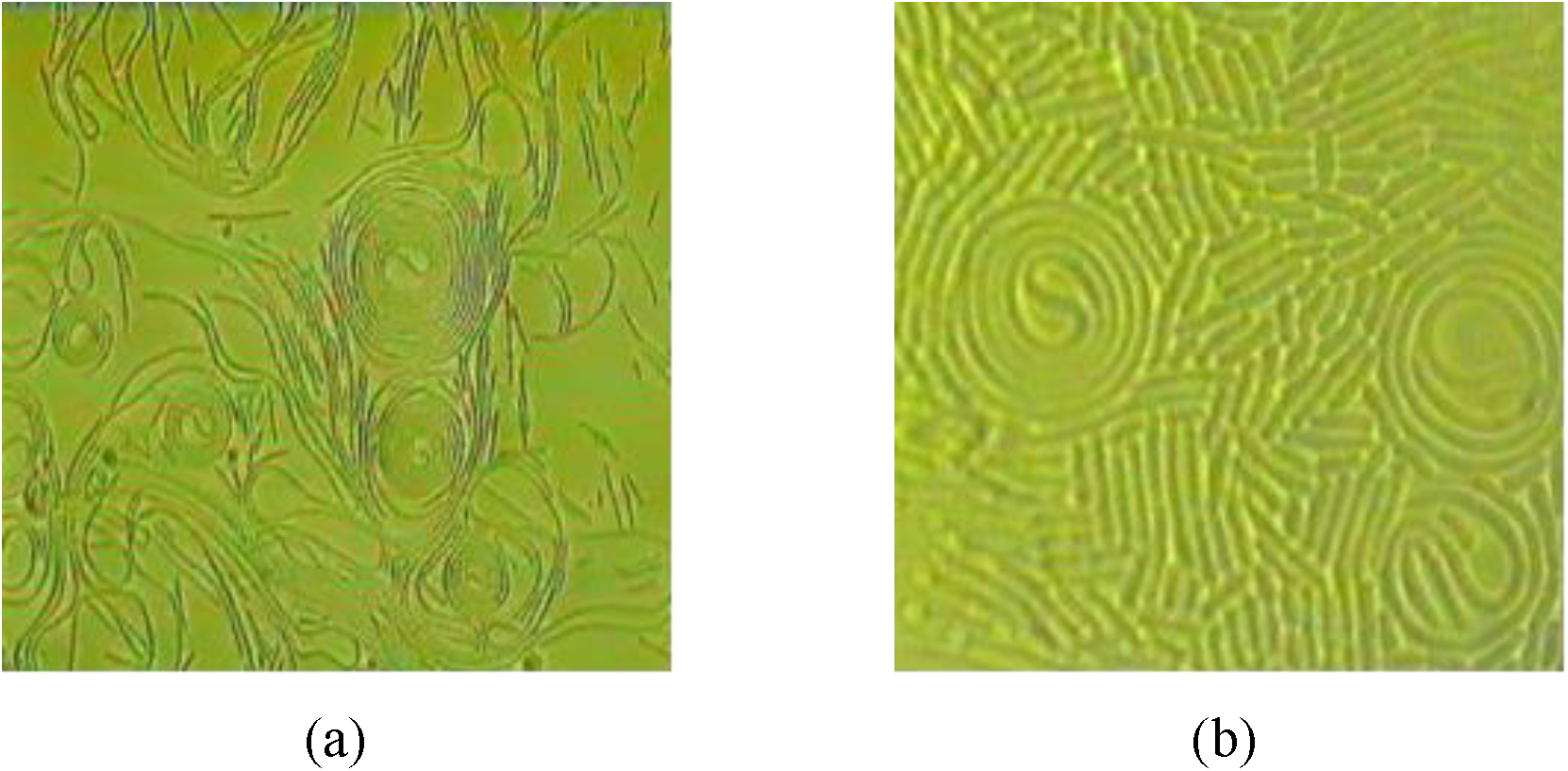
Under microscope observed phenomenon three bio-spins with diverse forms rotation movement were observed. The left one is a typical structural bio-spin. In its center there is an S-shaped hover as the supporting structure. During the process of rotation, both ends of S constantly change respective size and space to adjust the integral structure stability. While the structure of the top-right bio-spin is relatively simple, it constantly changes the size of the inter space to adjust the stability. The bottom-right one shows that the tendency of stability has already decreased. The “cryptic growth cell” rotation dynamic trend was observed in a visual field. The characteristics appeared every bio-spin rotation diverse direction as randomness whereas the speed almost uniformity which shows the power stability of the active movement.

#### 2.5 Filament cell powerful coupled oscillation

The powerful coupled oscillation of cryptic growth cell population as filaments could be more beneficial movement. Such as micro-turbulence phenomenon incubated at the temperature 37°C in 40 minutes the high speed rotation movement came into being. A sequentiality random dynamic process while at temperature 21°C form the chaotic turbulence. Mixed with the bio-spins impact spindrift occasionally form high rotation. It acted out huge kinematic characteristics. In this article also as the cryptic growth cell of proteus mirabilis strain produce a microcosmic turbulence phenomenon. It is the bacterial “cryptic growth cell”of proteus mirabilis incubated at the temperature 37°C in 40 minutes can appear high speed rotation, appearing rule and sequentiality dynamic process; while at temperature 21°C appeared the chaotic process of turbulence. The phenomenon that was observed under the microscope seemed like the famous Qiantang tide in southeast of China. Mixed with the bio-spin impactive spindrift which unthreaded occasionally with a high rotation speed, it acted out huge kinematic characteristics, as shown in Fig.8.

**Fig.8.**
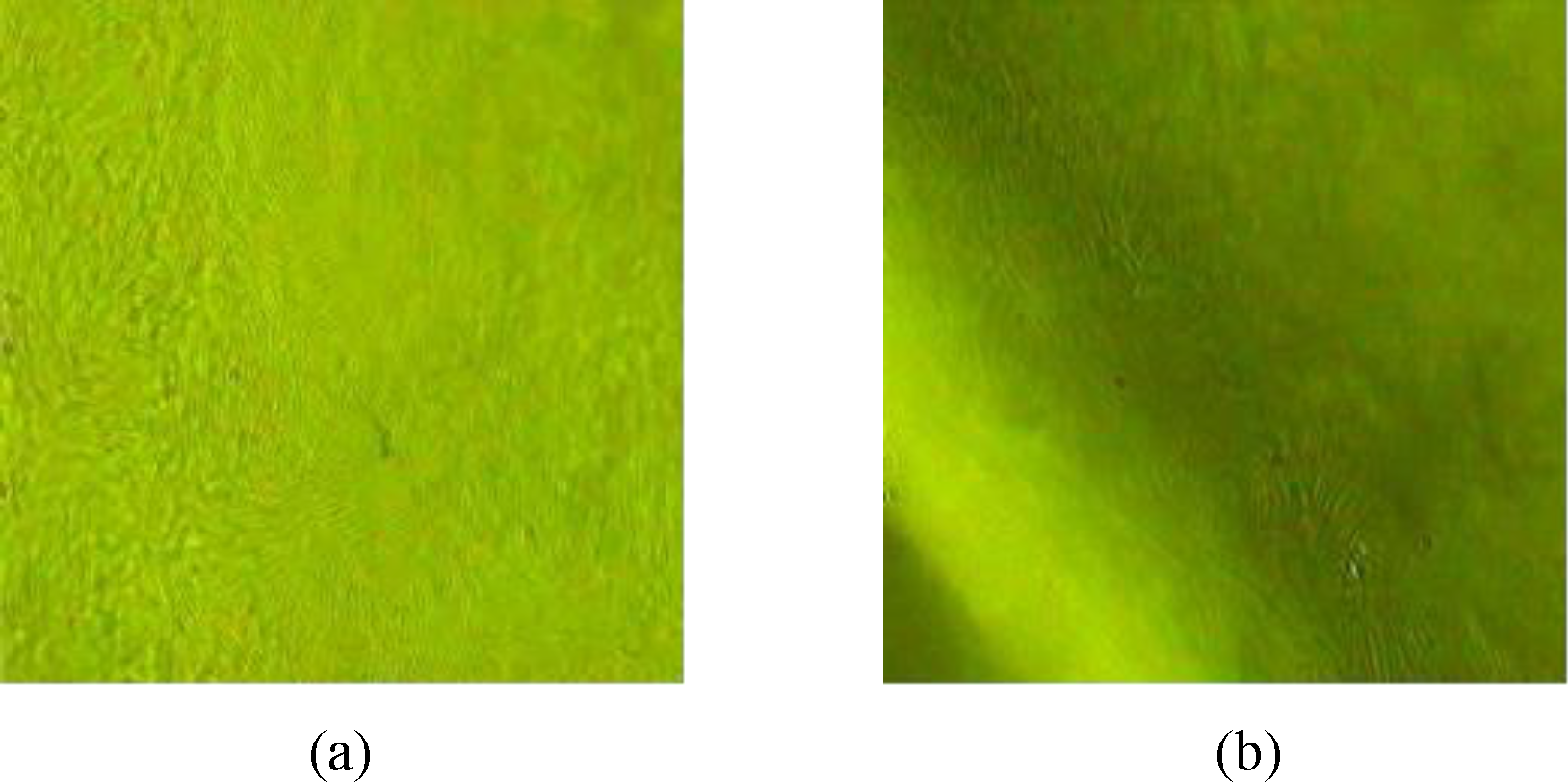
“cryptic growth cell” group turbulence dynamics.

The Fig above (a) showed the flowing-water-like phenomenon which is formed by “cryptic growth cell” group. (b) The turbulence dynamic process like the river surges up and down. Biowave basic experiment systemic observed the bacterial cryptic growth cell as filament population germinated a special motion process. About life organism generate the coupled oscillation dynamic process as well as along with the turbulence arises in living tissues are systematically described. It always is called as under microscope turbulence [4] that derived from the coupled oscillation dynamic whereas it opens up new possibilities to understand the new field of dynamic change in vivo. This is clearly paving the way for the opening up of the living organisms system to promote the integration of biomedical science into the scientific community. The biowave exploration trend has been clearly marked by the systematic entry of the system into sequential dynamic negative entropy flow to prevent and control cancer. The practice of medicine is realized in practice process.

### 3. Establishment of non-tumor mouse experimental model

In biowave basic experiment shows that both biotic communities from reproduce of the same species were continuously alternating. These phenomena may belong to the manifestations of homogeneity and heterogeneity in life organism. So, the transform pattern affects the whole of growth system. The basic model of biowave experiment has confirmed the effects of alternating on the species continuation. Similar situations often happen in terms of oncogenesis. This is also very useful for understanding the process of biological development. Although among these field there are certain differences in persistent period. In the course of life history was actually not completely independent of each other unless a new life begins. It is always used to establish transplanted and metastatic laboratory models.

#### 3.1 Natural non-tumor” mouse phenomenon

In long-term studies by a chance an unusually small tumor was observed next to a tumor that was locally inoculated with B 16 cell line. The cells of the small tumor were inoculated healthy mice. The results in vivo tumorigenesis ability was negative that is called as “natural non-tumor phenomenon”. This was quite unexpected that showed in addition to can’t form tumor in healthy mouse, the most notably was the experimental mouse rapidly became extremely active and healthy. So as to the healthy mouse could be endowed with the trait of non-tumor mice. Analysis suggests, the internal environment of life organism is the basic condition of tumor occurrence or anti-tumor ability to form. Maybe it lies in the selection of the heterogeneity composition of tumor cells rather than the whole cell structure in a special environment. And it could facilitate natural non-tumor mouse comes up. So, further study should focus on the hypoxic dynamics of internal environment. The first should start the experimental induce the formation of the EP oncocyte strain.

#### 3.2 EP oncocyte line

The EP oncocyte is a tumor cell which is thought to be the key inducer in directing the site-specific mutation of mouse lymph node cells. Mainly adopts physical and chemical approach to continuous stimulation the local skin of mouse submandibular lymph nodes. The surface specific antigen markers of lymphocytes gradually fade away as fig.9. Then it was screened by cell culture furtherly. Such screened cell was designed to external point induce directing site-specific of lymph node cytometaplasia called as “external point mutagenic cell strain” referred to as EP oncocyte. That is a key evidence of hypoxic change of internal environment in tissue structure. In experimental process showed that the tissue structure trait of lymph node under inflammation influence was similar with of tumor ecological niche.

**Fig.9.**
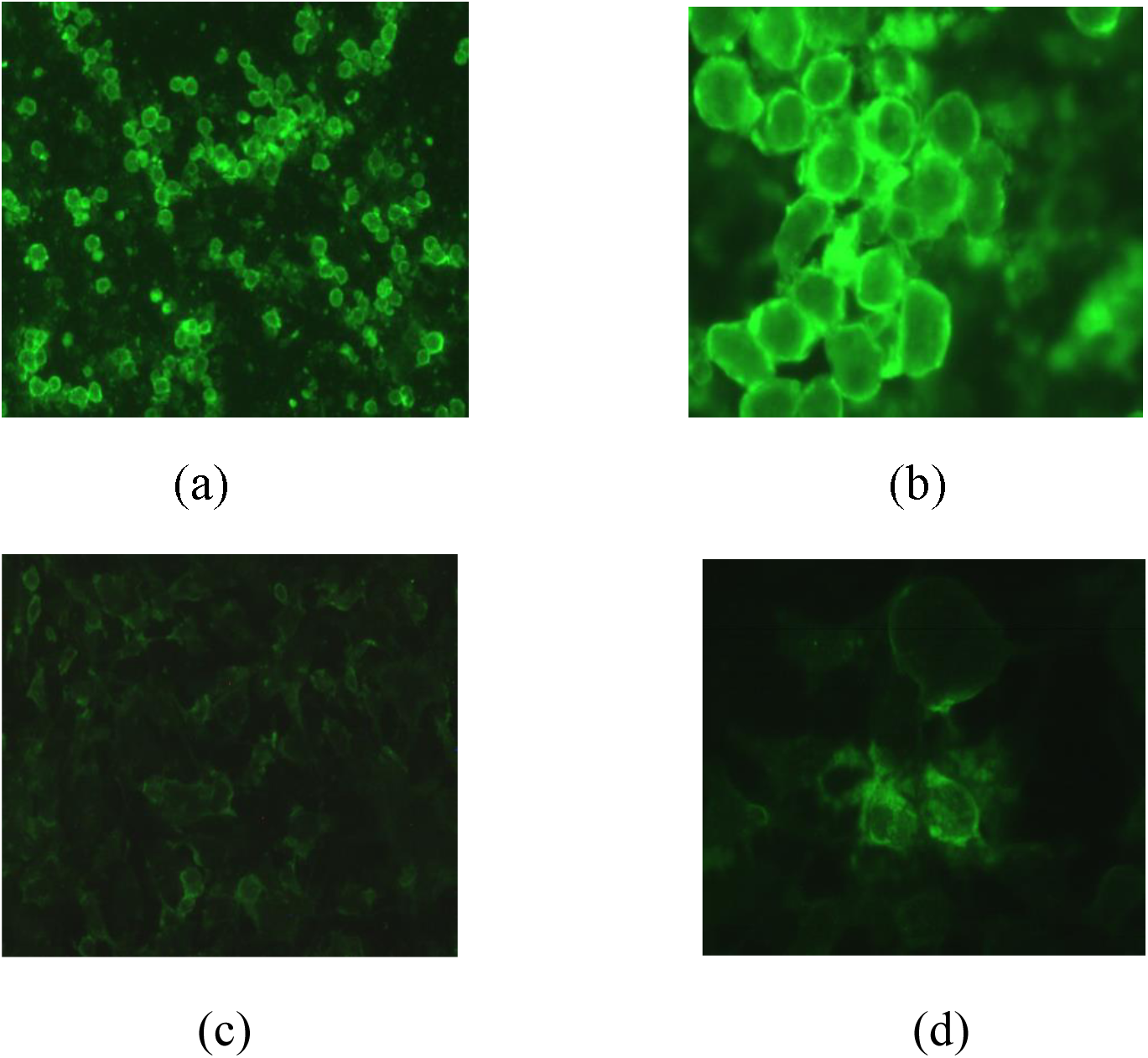
showed the change process of the physiological landmark molecule on the lymph node cell membrane surface.

The fig.9 above (a) shows the lymph node cells test stain with the specific immunofluorescence labeled. (b) The lymph node cell induced 10 days. (c) Induced time had reached 15 days. (d) Had induced all the cells was became tumor cells.

#### 3.3 EP non-tumor mouse model

EP oncocyte emergence lays the foundation for a wider field of biowave anticancer research. The operation method took after the emergence of natural non tumor mouse phenomenon to establish the EP oncocyte non-tumor mouse model. The standard of preparation of EP oncocyte non-tumor mouse models should include the following aspects: cancer nest of specimen tissue criteria; Lot of slender tubular has been collected as a cytology index; Cell composition extraction and treatment should be include grinding into homogenate and heating inactivated; then precipitation and collect-supernatant liquor as operational procedures. The non-tumor mouse model constructed with EP oncocyte is called as EP non-tumor mouse model. From a scientific point of view that the classical experiment of this aspect rests with the establishment the experimental model of the murine melanoma B 16 cell non-tumor mouse.

#### 3.4 The non-tumor mouse model

Oncology laboratories have long been used to study B 16 cell clones. On the basis of EP non-tumor mouse model the build method is also suitable B16 cell cline. So the latter is simply called non-tumor mouse model.

#### 3.5 Cross resistance tumorigenesis

With both non-tumor mouse model established were certain to bring about the biowave anti-cancer deep-going study. The most importantly, these models showed tumorigenic cross resistance. That is in both non-tumor mouse models, each could resist tumorigenesis of EP oncocyte and B16 cells strains. In the process of building the two non-tumors mouse models the preparations necessarily were respectively prepared. It’s an idea clearly worth factor that closely related to their matrix trait able to withstand the processing operation of heating inactivation, as well as can through microfiltration membrane. After trial and error the results show that cross anticancer efficacy of these preparations lies in the same mechanism of efficacy. Especially the two microstructure of a single physical heat resistance is basically the same. It is inferred, this signifies the new way of cancer prevention and control lie in the nonspecific single physical properties of the carbohydrate components of tumor cells. And different tumor cells all contain the same substance. Its material properties have been touched upon in tumor preparation of constructing model.

### 4. RNA constituent prepared and standard determined

The basic standards of the experimental model of non-tumor mouse model must be strictly referenced. In addition to model resistance to tumor formation the RNA component structure standard and experimental analysis are mainly involved. Furtherly enter human cancer cells experiments should be direct beginning from matrix of the preparation.

#### 4.1 Observation of preparation matrix under the microscope

In allusion to the micro-particles released by tumor cells on the surface of BS solid medium and incubate at 37°C. Observe under microscope shows that a large number of slender-shaped structures distribute in a form of intertexture, which were accompanied by a mixture of micro-particles with linear parallelism as can be seen in Fig.10.

**Fig.10.**
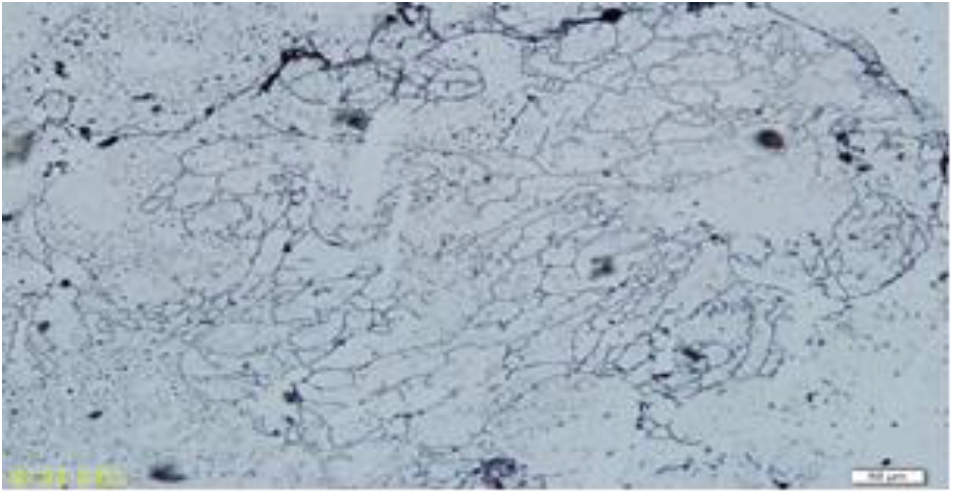
Observation of the preparation under optical microscope.

It shows that the elementary composition of theRNA 1 2 3 preparation matrix mainly includes slender-shaped structures and microcosmic particles in different sizes. The diameters of the bigger particles are around 0.1μm. It can be seen that most of the elementary composition appears with beaded-like array. It can be seen that most of the elementary composition appears with beaded-like array.

#### 4.2 AFM Scan the microcosmic particles under DFM mode

As for the elementary composition of RNA 1 2 3, on the basis of the optical microscope observation, further scan the slender tube shaped structures and the subtle feature of microcosmic particles, as well as the relevance between the above two under the DFM mode of atomic force microscope (AFM). The results are shown in Figs. 11 and 12. The scanning result shows, the elementary composition of the preparation matrix has been confirmed. It is speculated that, the relevance of the above two probably is that the microcosmic particles are stored in the slender tubes of the living cells. The sequential vibration of microcosmic particle population promotes the slender tubes to behave the similar vibration to the coupled oscillation process of “cryptic growth cell” in biowave growth.

**Fig.11.**
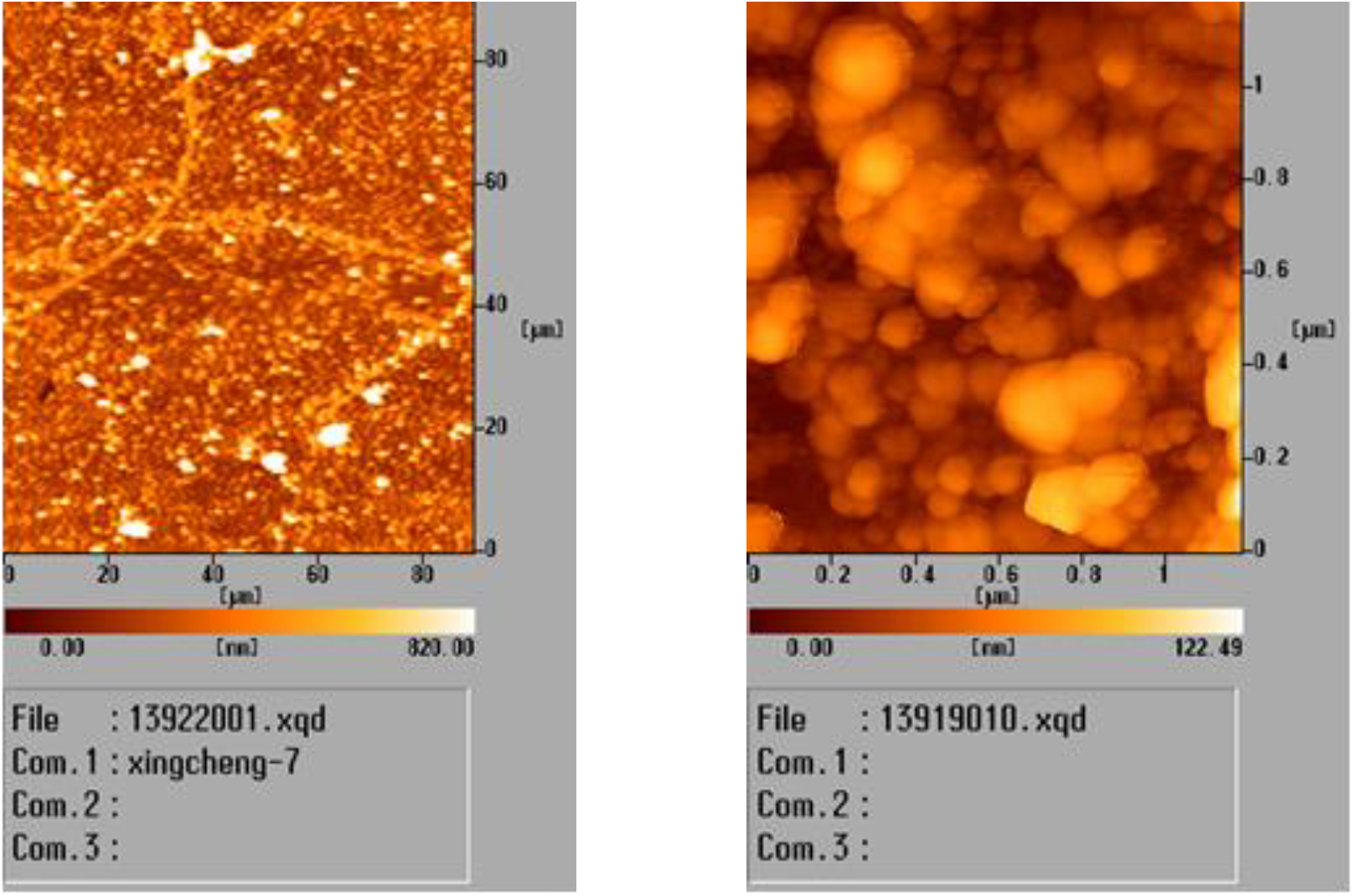
DFM Scanning in the scope of 80μm (left) and 1μm (right)

**Fig.12.**
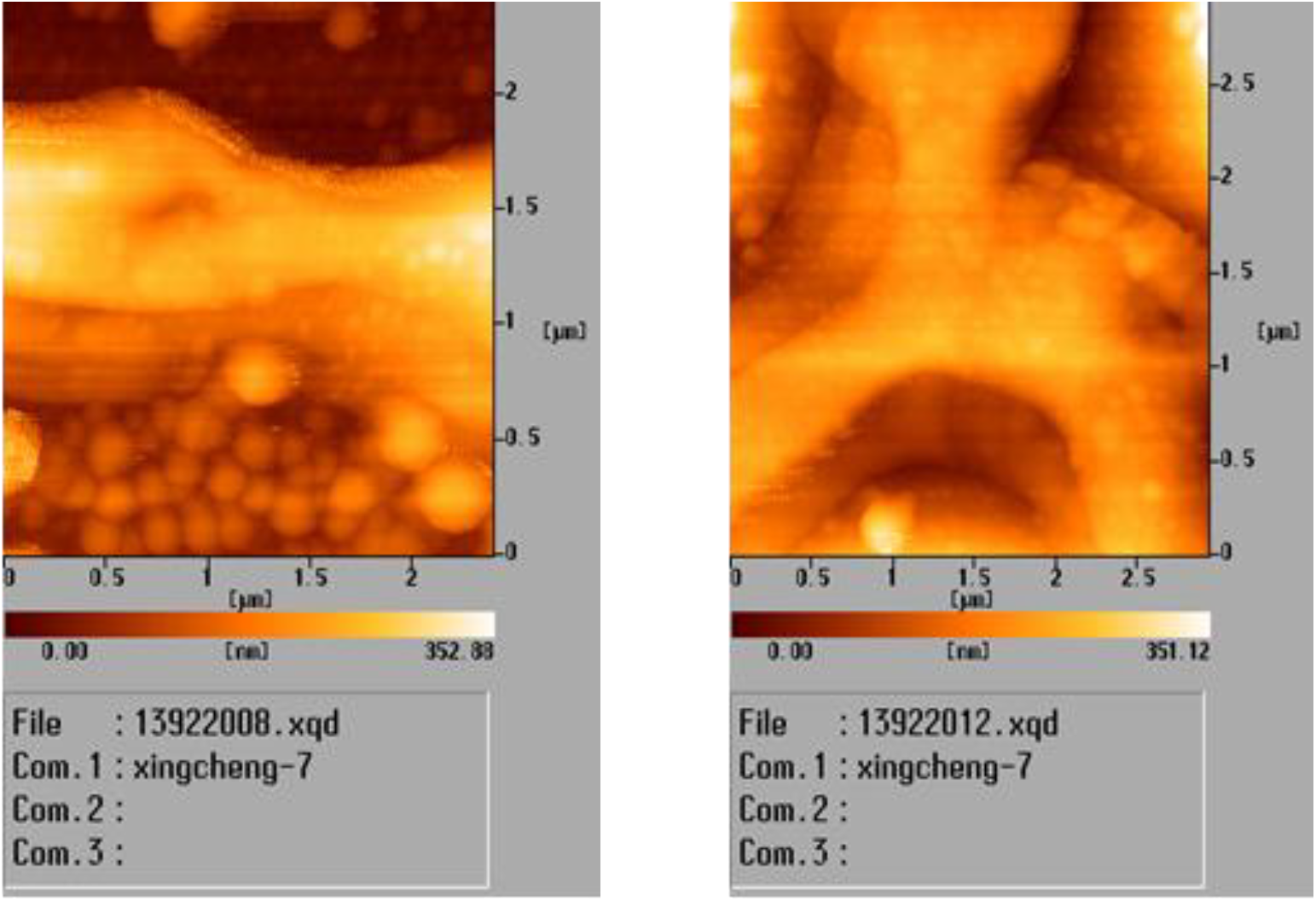
DFM Scanning the shape of slender tube and the micro-particles.

The Fig. above shows out plenty of accumulated intensive microcosmic particles. In general, the size of microcosmic particles is relatively homogeneous. It also shows out the slender shaped structure which is crookedly folded. It shows the thickness of the slender shaped structure is relatively consistent with the bigger microcosmic particle. The high magnification image shows that the sizes of microcosmic particles are obviously heterogeneous. Diameter of the bigger one is 100-200 nm, while diameter of the smaller one is 30-60 nm.

The scanning phenomenon of slender tube shaped structure as mentioned in Fig.10. The accumulation of microcosmic particles can be seen under the tube at the depth of about 100nm (lef). Therefore, the picture shows the connection between slender the tube shaped structure and microcosmic particles on the aspect of deep leveled change. It seems that the tube shaped structure cracked and the microcosmic particles discharged out of the tube. According to the result of observation, it is possible that there are microcosmic particles being stored inside the slender tube. The image at right shows that the standing out circular structure is higher than the surface of slender tube. The height is around 100-200 nm. It can reflect the relevance between slender tube and microcosmic particles on one side. Similarly, it shows that microcosmic particles with the height of around 100 nm faintly appear under the position of the tube. There exists a microcosmic particle with the height of 300 nm approached to the deep crack of the slender tube.

#### 4.3 Nucleotide gel electrophoresis identification have already done

Aqueous solution specimens of tumor cells RNA particle via boiling processing is divided into1, 2, 3, 4 parts. Then these samples were disposed with enzyme digestion process respectively. The results represent the RNA existence only. No DNA was observed. It is shown in Fig.13.

**Fig.13.**
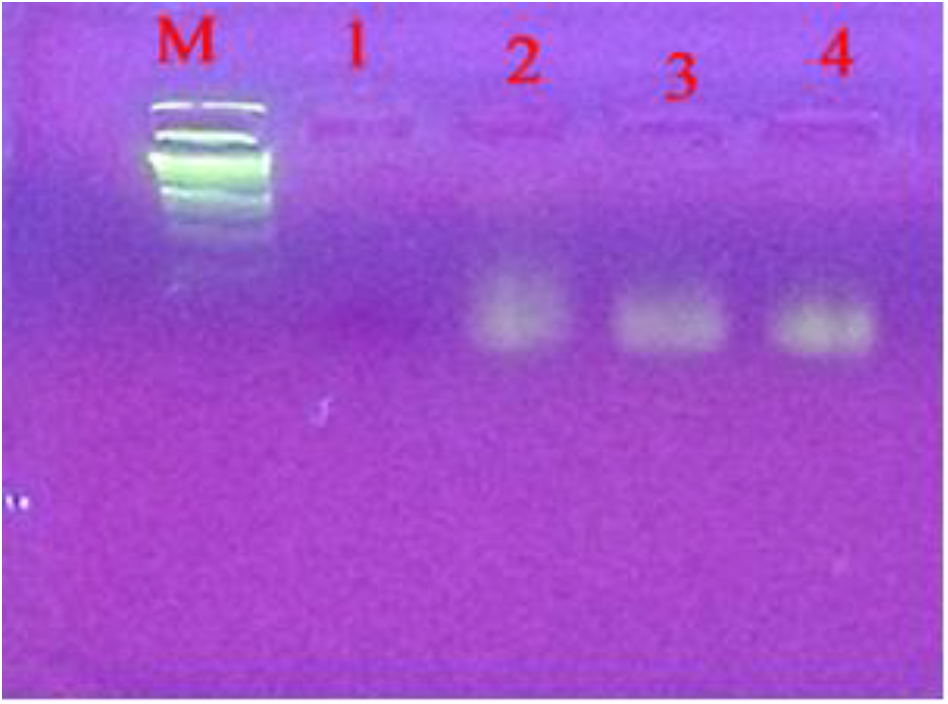
Nucleotides gel electrophoresis.

The specimen of the RNA preparation was the aqueous solution of grinding from tumor cell. After boiling the test specimen was divided into 1,2,3,4 samples which were managed respectively. The marked by DNA Marker, No. 1 sample as test that with RNA enzyme digestion treatment. No. 2,3,4 as the control test under different conditions have not get any treated included RNA enzyme managed. Since these control only appearing RNA bends it is proved the existence of RNA. The analysis suggests that the RNA constituent can form infinite kinetic physical strength by means of the vitality in life organism. This is an infinite power that strengthens the physique that is nonspecific anti-cancer efficacy principles.

### 5. Preclinical trial indicators

To study the RNA vaccine for preclinical trial the experiment shifted to the indicators of preparation. The preclinical trial research should be direct beginning from the basic standards of the RNA. Especially the experimental models of non-tumor mouse must be as clinical set of rules to follow.

#### 5.1 Evaluation of preliminary effect

It includes the pharmaceutical indicators that to determine water evaporation index. It is as to adjudge the micro-turbulence index of the RNA preparation.

#### 5.2 The effectiveness evaluation of the RNA constituent

Based on the effectiveness of experimental models of non-tumor mice it should be quantitative expression efficiency of cancer by biowave effect time uncompressing assay. The prevention and control as well as the time to take effect and continue need to be clearly shown.

#### 5.3 The safety evaluation

It mainly considers the allogenic rejection reaction and allergic reaction test criterion. Fortunately, it has been destroyed by inactivating preparation processing.

## DISCUSSION

About the action of tumor cell RNA studies begin with the concept of uncertainty. Acquaint oneself with the biological phase variation phenomenon of single bacillus population. It from the bacterial vegetative cell turns into filaments as “cryptic growth cell” that the innumerable filament could cause incessant dynamic process of coupled oscillation. The violent turbulence under microscope shows the infinite imagination space in front of biowave researchers. Subsequent research went on until it can be known to be used in anticancer. With the help of vitality in life organisms the coupled oscillation of this single physical structure can persistently arise the coupled oscillation of this single physical structure can persistently arise as if a biological behavior. To pay back the whole body system maintaining negative entropy flow sequence dynamic process. Thus, the apparent result of life organism system openness is quite different with the concepts involved in dissipative structure theory. The complexity of living system is based on the formation of a nonlinear relationship between the single physical RNA constituent and the tissue structure vitality of living organisms. The former has the restriction of dissipation process while the latter will be walk with life. This point has been preliminarily verified in RNA cancer vaccine mice experiment model. At present there are two aspects can be clearly clarified. The first as its entry-into-force conditions must be in the homogeneity of the primary tissues in the whole body. The next is the rapid, extensive and long - term effectiveness rely on the tissue embryonization efficacy derived also from the RNA vaccine. It can play biowave dynamic rejection heterogeneity powerful role during its selection process. The RNA composition as a single physical structure still can show biological behavior that the special dynamic effect may be involved some new function. It including the ability to choose and correct deviate from physiology. The systematic studies have focused on the theoretical basis of RNA vaccine anticancer action mechanism. The principle of its efficacy has been clarified that show the specific action of the single physical power can convert to biological behavior. In the process one is called biowave life vitality always can bring up specific procedure of tissue embryonization. It deal with the species stability that keeping life vigor generalized explanatory.

## CONCLUSION

As for the experiment of RNA anticancer argumentation mainly focuses on the RNA component consist of inactivated structure can produce singly appropriate biological activity effect. The condition must accompany the life organic whole life movement vitality. Systematic study results show that the basic working modes of RNA element are to give rise to and take part in coupled oscillation in cytoplasma with biochemical metabolism. So, it is closely connected biological couple oscillation. About RNA component has been mainly referred to the relationship between inactivated beaded structures and life organism physiology vitality. The former are wrapped around mucosal tissue that can form biological oscillator. The latter derives from internal and external environment communication that can produce trigger factors. Finally the force source favors oscillators sequence to form collective rhythm and coupled oscillation occurs. The coupled oscillation shows the response to dynamic variations of environment and reflects continuous communication process. More precisely, in this process the first defines the difference of nonliving structure and living behavior could fuse together in the random response of chaotic oscillators. The boundary between above two disappears in the biological coupled oscillation qualitatively. These have come into being since from the origin of life and gone through biological evolution in life organism. Thus, it clearly shows that fundamental capabilities of life organism induce physical properties. The basic research is from the beginning of exploration till to the determination of biowave oscillators. The oscillators in coupled oscillation take part in the dynamic process of biochemical metabolism in life organism. Finally, the research pursues microcosmic motion of life organism. The pursuers have eventually realized that the sequential motion of biowave oscillator is exclusively initiative force of life organism apply work. The climate is extreme in the northeast of China that is suitable for the research about life organism and nature environment. At last, the mechanism of maintaining communication between life organism and nature has been clarified. The new concept of life oriented temperature difference convection. The latest research advance reveals fundamental metabolic pathway of life organism acquiring vitality. The vitality can be regarded as to power with biology apply work. Biowave theory shows the process of physiological apply work necessarily comes with dynamic sequence of negative entropy flow in whole body. On this basis, life five-step chain mechanism of life organism vitality linking with system openness has been clarified. From this, it reveals that special apply work comes from life oriented temperature difference convection. New medical technique will be established on the basis of causal circle principle. Laws of RNA will pioneer new areas of medicine to solve medical difficult problems.

---

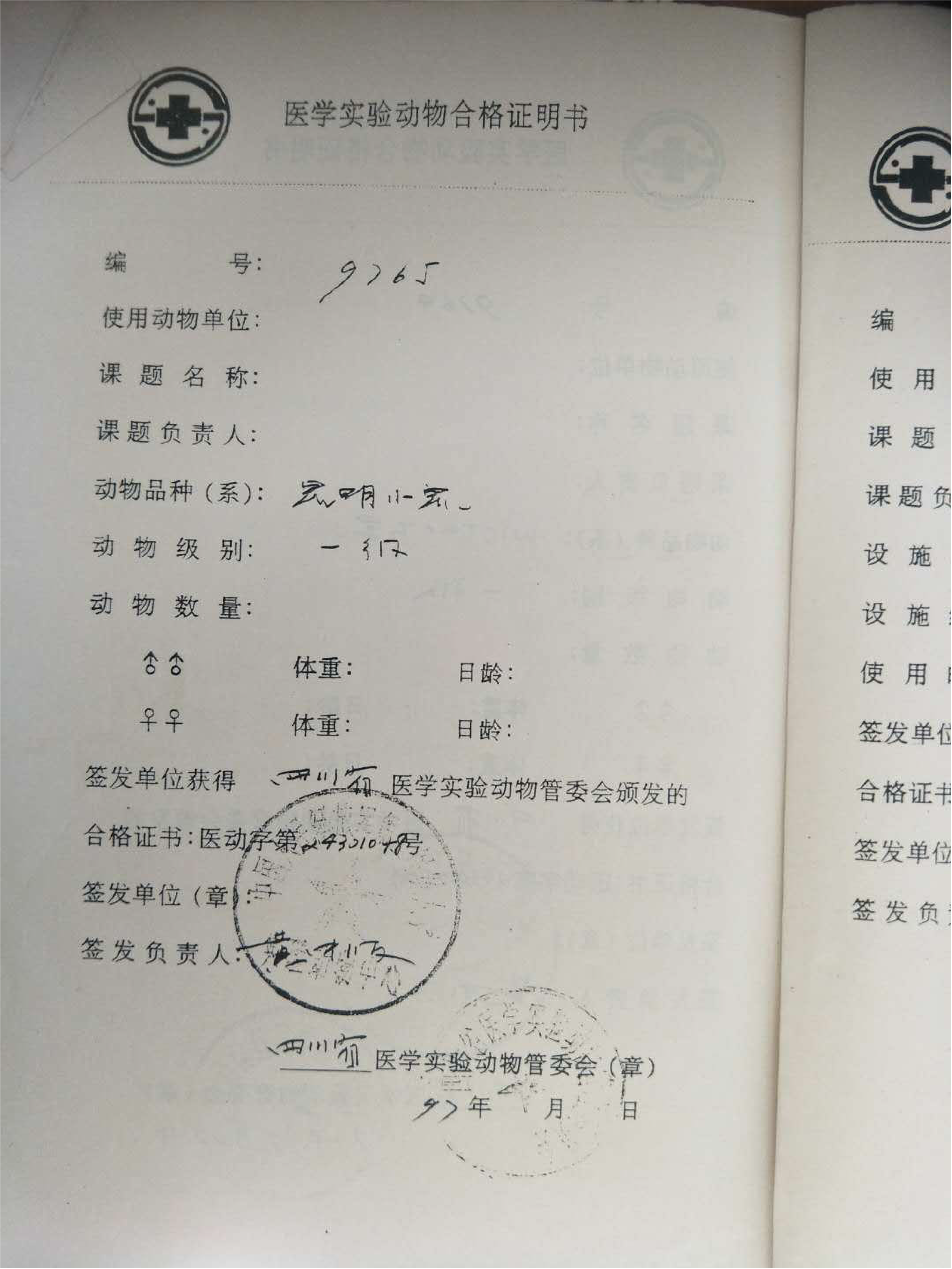

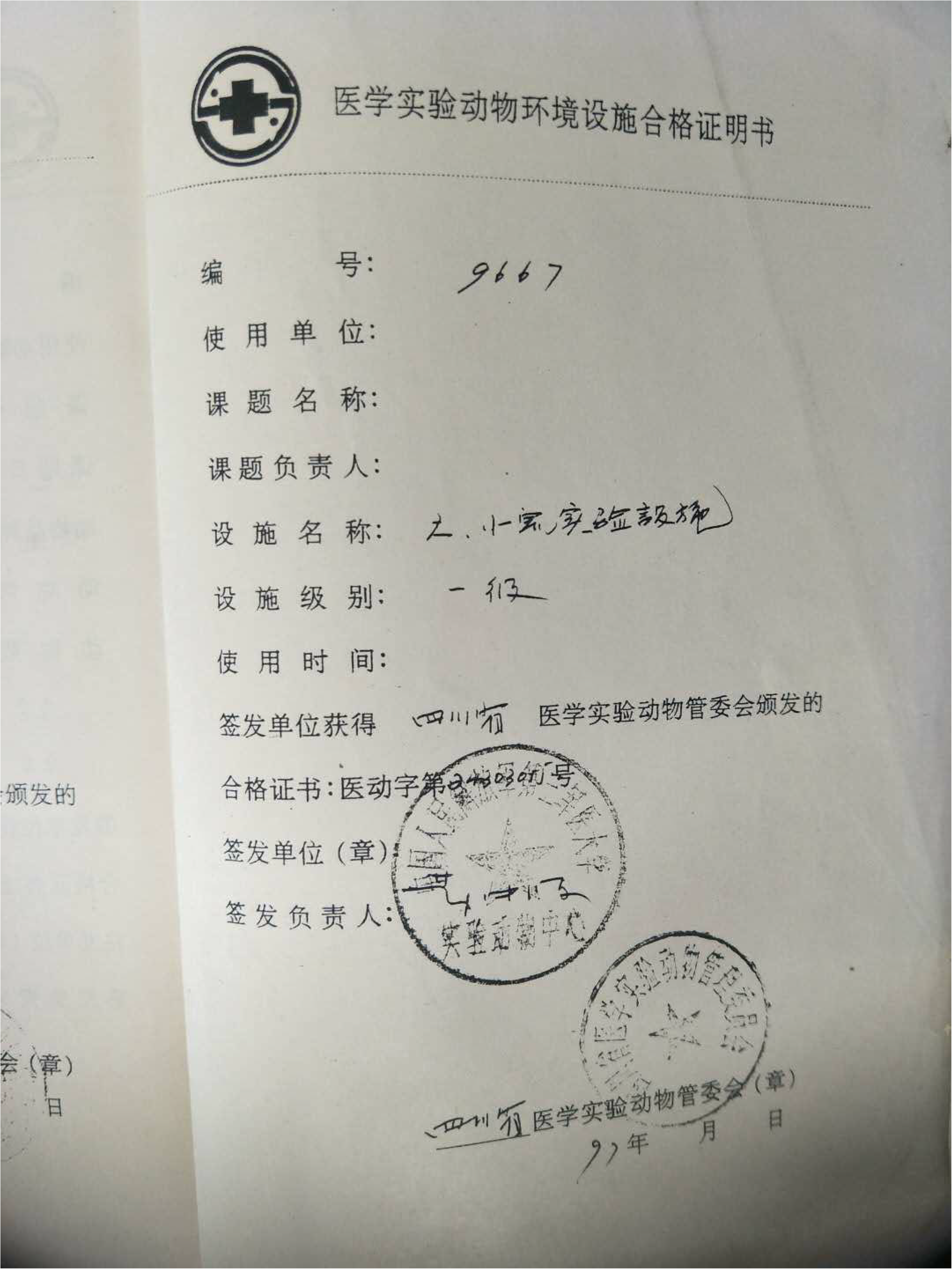

